# Scale-dependent effects of geography, host ecology, and host genetics on diversity of a stickleback parasite metacommunity

**DOI:** 10.1101/677161

**Authors:** Daniel I. Bolnick, Emlyn J. Resetarits, Kimberly Ballare, Yoel E. Stuart, William E. Stutz

## Abstract

Many metacommunities are distributed across habitat patches that are themselves aggregated into groups. Perhaps the clearest example of this nested metacommunity structure comes from multi-species parasite assemblages, which occupy individual hosts that are aggregated into host populations. At both spatial scales, we expect parasite community diversity in a given patch (either individual host or population) to depend on patch characteristics that affect colonization rates and species sorting. But, are these patch effects consistent across spatial scales? Or, do different processes govern the distribution of parasite community diversity among individual hosts, versus among host patches? To answer these questions, we document the distribution of parasite richness among host individuals and among populations in a metapopulation of threespine stickleback (*Gasterosteus aculeatus*). We find some host traits (host size, gape width) are associated with increased parasite richness at both spatial scales. Other patch characteristics affect parasite richness only among individuals (sex), or among populations (lake size, lake area, elevation, and population mean heterozygosity). These results demonstrate that some rules governing parasite richness in this metacommunity are shared across scales, while others are scale-specific.

## Introduction

A long-standing question in ecology is, “why are some communities more diverse than others?” To address this question, biologists have sought to identify and quantify the biotic and abiotic factors that affect community diversity, and which differ across a landscape. A clear lesson from such work is that the salient factors depend on the spatial scale being considered (Cottenie 2005; Chave 2013; Meynard *et al* 2013; Chase and Leibold 2017). For instance, ecosystem productivity is often negatively associated with community diversity at small spatial scales, but the trend can be reversed at large scales (Chase and Leibold 2002; McBride *et al* 2014). Similarly, here we show that factors affecting parasite metacommunity diversity change depending on the spatial scale at which the communities are defined.

Parasites make up a large proportion of biodiversity (Poulin and Morand 2000) and have major implications for community dynamics (Lafferty *et al*. 2008), conservation (Thompson *et al*. 2010), and human health. Yet, the processes structuring variation in parasite diversity remain poorly understood (Mihaljevic 2010; Presley 2011; Moore and Borer 2012; Richgels *et al*. 2013; Dallas and Presley 2014; Borer *et al*. 2016). To better understand the rules governing the distribution of parasite richness, parasite ecologists are increasingly drawing on metacommunity theory (Grenfell and Harwood 1997; Ebert *et al*. 2001; Leibold *et al*. 2004; Seabloom *et al*. 2015). For parasites, each individual host represents a transient habitat patch that often supports a multi-species parasite community (an ‘infracommunity’ per disease ecologists). A single host population can thus be viewed as a parasite metacommunity (a ‘component community’). Or, we can view each discrete host population as a habitat patch for parasites, and a collection of host populations form the metacommunity. Parasites therefore form a spatially nested metacommunity, whose diversity and composition varies among host individuals within host populations, and among host populations (Borer *et al*. 2016). Most research on the ecological processes governing parasite diversity have emphasized one of these spatial scales (Poulin 1997; Vidal-Martínez and Poulin 2003; Poulin 2007). There is therefore a need for more studies that bridge both spatial scales to ask whether the same processes regulate parasite diversity at both the host individual, and host population, scale?

Here, we examine scale-dependent effects on parasite richness (α-diversity), using a host metapopulation of threespine stickleback (*Gasterosteus aculeatus*) on Vancouver Island. Parasite infection prevalence in stickleback is known to covary with individual hosts’ diet, morphology, sex, and immune genotype (Reimchen and Nosil 2001; Stutz *et al*. 2014; Stutz and Bolnick 2017). Parasite diversity and community composition also differ among stickleback populations (MacColl 2009; Eizaguirre *et al*. 2011; Poulin *et al*. 2011; Stutz *et al*. 2014, 2015), due to among-population differences in host immune genes (Matthews *et al*. 2010a; Eizaguirre *et al*. 2011; Stutz and Bolnick 2017), diet (Matthews *et al*. 2010a), and abiotic conditions (Simmonds and Barber 2015). These many studies of infection in stickleback yield inconsistent conclusions, perhaps because factors regulating infection have scale-specific effects. To evaluate this possibility, we considered five broadly applicable predictions concerning the host-individual and host-population ‘patch’ traits that structure parasite local community diversity. We focus on parasite community richness here, a companion paper (Bolnick et al., 2019) provides complementary analyses focusing on variation in the abundance of particular parasite species.

### Prediction 1: Host diet has a scale-independent effect on parasite richness

In many animal species, co-occurring individuals actively prefer different prey, resources, or microhabitats (Bolnick *et al*. 2003). Host populations also diverge in diet and resource use, often along the same ecological axes as the individual-level diet variation (Schluter and McPhail 1992, Matthews *et al* 2010). Because many parasites are trophically transmitted (Wilson *et al*. 1996; Johnson *et al*. 2009; Cirtwill *et al*. 2016), host diet (composition or diversity) should correlate with parasite richness. This correlation should be consistent across spatial scales, assuming each parasite’s complex life cycle is relatively invariant across the metacommunity.

### Prediction 2: Host ecomorphology has a scale-independent effect on parasite richness

Diet variation often reflects variation in trophic morphology that affects prey capture and processing (Wainwright and Richard 1995). A corollary of Prediction 1 is that trophic morphology should be correlated with parasite richness. Indeed, morphology might be a more reliable covariate of parasite richness because diet measurement is noisy and often restricted to short time scales. If the diet-morphology relationship is consistent across host populations, morphology effects might therefore also act similarly across spatial scales.

### Prediction 3: Host heterozygosity has a scale-independent, negative effect on parasite richness

Parasite diversity should be inversely related to the host’s ability to recognize and eliminate various parasites (given equal exposure risk). Such host immune competence can be related to genetic diversity at particular loci (e.g., MHC; Wegner *et al*. 2003; Kalbe *et al*. 2009; Oliver and Piertney 2009), but immunity will often depend on the collective action of many loci. Thus, another measure of immunogenetic diversity might be genome-wide genetic heterozygosity (Coltman *et al*. 1999; Arkush *et al*. 2002; Whitehorn *et al*. 2011). We predict that genome-wide heterozygosity is negatively related to parasite richness among host individuals and among populations. Alternatively, parasite richness might be controlled by host immune genotypes at specific loci that affect resistance to many taxa at once. If such loci evolved in parallel across replicate populations, we should find richness associated with certain SNPs’ genotypes, at both spatial scales.

### Prediction 4: Host sexual dimorphism contributes only to within-population variation in parasite richness

In many species, sexes differ in feeding ecology (Shine 1991) and immunity (Zuk 1996; Nunn *et al* 2009), which should influence individual-level parasite richness. Because host populations have a roughly equal sex ratio, we do not predict that dimorphism will influence among-population variation in parasite richness.

### Prediction 5: Habitat contributes to among-population differences in parasite richness

Elevation, lake size, and distance from the ocean are geographic features that should be experienced by all individuals living in a given site. Therefore, they should contribute only to among-population differences in parasite richness (Ebert *et al*. 2001; Anderson *et al*. 2010; Johnson and Thieltges 2010; Richgels *et al*. 2013; Johnson *et al*. 2016).

## Methods

### Collection

In June 2009 we collected threespine stickleback from 46 sites on Vancouver Island, British Columbia, Canada, in the historical lands of the Kwakwaka’wakw First Nations. Our sample sites included 5 estuaries with anadromous fish, and 33 lakes and 8 streams from 9 watersheds (Table S1, Fig. S1). At each site, we placed unbaited 0.5-cm gauge wire minnow traps along ∼200m of shoreline in 0.5-3m deep water until we took 60-100 fish per site (Table S1). Fish were euthanized in MS-222 and preserved in 10% buffered formalin after saving a fin clip in ethanol for DNA. Specimens were later rinsed and stored in 70% isopropyl alcohol after staining with Alizarin Red (brand). Collection and animal handling were approved by the University of Texas IACUC (07-032201) and a Scientific Fish Collection Permit from the Ministry of the Environment of British Columbia (NA07-32612).

### Parasite diversity

We scanned each fish under a dissection stereomicroscope to count and identify macroparasites (helminths, crustaceans, mollusks, and microsporidia) to the lowest feasible taxonomic unit, following Stutz and Bolnick (2017). We focus on two parasite diversity statistics (Figure S2): (1) the number of uniquely identifiable parasite taxa for each fish, α*_ij_*, i.e., parasite richness in individual host *i* from host population *j*; (2) the average per-individual parasite richness, 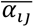, for all fish in population *j*. (Details on parasite taxonomic composition and abundance are reported in the companion to this paper (Bolnick *et al,* submitted).)

### Morphology

Before dissection, we weighed all fish within 0.01g and used digital calipers to measure standard length, body depth, and body width at the pectoral fins (mm). For a random subset of 30 individuals per population, we measured three trophic traits—gape width, gill raker number, and longest gill raker length—and averaged left- and right-side armor plate number. We sexed all individuals by visually inspecting gonads. Linear measurements were log transformed and size-standardized by regression on log standard length. We calculated trait means per population for population-scale analyses.

### Diet

For a random subset of 28 populations, we analyzed fish stomach contents in the same 30 fish used for ecomorphology, recording presence of prey to the lowest feasible taxonomic level to calculate the number of observed prey taxa per fish (prey richness). Stickleback individuals and populations vary in their relative consumption of benthic invertebrates versus pelagic zooplankton (Lavin and McPhail 1986), so we use the proportion of benthic prey taxa as another diet metric. This measure was highly correlated with the first axis of a non-metric multidimensional scaling (NMDS) analysis using fish stomach contents (per the Jaccard index in the R package *vegan;* Table S2). Several studies have confirmed that stomach contents are correlated with direct observations of feeding behavior, with long-term measures of diet using stable isotopes, and with trophic morphology (Matthews *et al*. 2010b; Snowberg *et al*. 2015). *Genetic diversity.* With DNA from a subsample of 12 fish per population for four marine sites, 31 lakes, and six streams, we used ddRADseq (Peterson *et al*. 2012) to obtain genotypes for 175,350 SNPs in 336 fish (mean 107,698 SNPs/individual). The protocol and bioinformatic pipeline are described in (Stuart *et al*. 2017). We calculated genome-wide mean heterozygosity (*H_ij_*) for each fish *i* in population *j*, and mean heterozygosity for each population (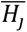).

### Statistical Analyses

All statistical models were implemented in R version 3.5.2 (see Table 1 for full list). Our primary goal in this study is to understand why host individuals and host populations differ in parasite α diversity. To understand the relative effects of population and watershed identity, we fit a Poisson General Linear Model (GLM) evaluating how per-fish parasite richness varies as a function of host population nested within watershed (both random effects, see model M1 in Table 1). For each source of variation, we calculated its effect size using *η*^2^.

**Table 1:** Models tested in this study. A summary of the statistical results of each model is provided in Supplemental Table S3. Blue rows focus on within-population variation and use individual fish as the level of replication (except M7), while yellow rows focus on sources of between-population variation and use population as the level of replication.

| Name | Model: | Does parasite richness depend on: |
| --- | --- | --- |
| M1: | $\text{logit}(\alpha_{ij}) \sim \lambda_j \mid \text{watershed}$ | host population, nested within watershed |
| M2: | $\text{logit}(\alpha_{ij}) \sim \lambda_j + \text{sex}_j + \text{length}_j$ | individual host sex and size |
| M3: | $\text{logit}(\alpha_{ij}) \sim \lambda_j + \text{sex}_j + \text{length}_j + \text{GRN}_j + \text{GRL}_j + \text{PN}_j + \text{GW}_j$ | individual host ecomorphology |
| M4: | $\text{logit}(\alpha_{ij}) \sim \lambda_j + \text{sex}_j + \text{length}_j + \text{NMDS1}_j + \text{NMDS2}_j + \text{DB}_j$ | individual host diet |
| M5: | $\text{logit}(\alpha_{ij}) \sim \lambda_j + \text{sex}_j + \text{length} + \text{SNP}_k$ | individual host genotype at SNP $k$ (GWAS) |
| M6: | $\text{logit}(\alpha_{ij}) \sim \lambda_j + \text{sex}_j + \text{length} + \text{Het}$ | individual host mean genome-wide heterozygosity |
| M7: | $t[\alpha_j] \sim t[\text{length}] + t[\text{GRL}] + t[\text{GW}] + t[\text{NMDS1}] + t[\text{NMDS2}]$ | sexual dimorphism in ecomorphology and diet |
| M8: | $\bar{\alpha}_{i,j} \sim \text{habitat} + \text{watershed}$ | habitat |
| M9: | $\bar{\alpha}_{i,j} \sim \log \text{lake size} + \text{elevation} + \text{ocean distance}$ | lake geography |
| M10: | $\bar{\alpha}_{i,j} \sim \text{mean length} + \text{mean GW} + \text{mean GRN} + \text{mean PN} + \text{mean GRL}$ | host population ecomorphology |
| M11: | $\bar{\alpha}_{i,j} \sim \text{mean NMDS1} + \text{mean NMDS2} + \text{mean DB}$ | host population diet |
| M12: | $\bar{\alpha}_{i,j} \sim \text{freq}(\text{SNP}_k) + \text{watershed}$ | host population allele frequency at SNP $k$ (GWAS) |
| M13: | $\bar{\alpha}_{i,j} \sim \text{mean Het}$ | host population mean heterozygosity |
| M14: | $\text{mean Het} \sim \log \text{lake size} + \text{elevation} + \text{ocean distance}$ | |

KEY: $\lambda_j$ : population-specific Poisson rate (random effect intercept); Subscript $j$ on any of the following traits indicates we fit a main effect and population random effect of a given trait (e.g., population by trait interactions); length: log transformed standard length; GRN: gill raker number; GRL: size-corrected gill raker length; PN: armor plate number; GW: size-corrected gape width; NMDS: diet ordination axes 1 and 2; DB: diet breadth; SNP: genotype at a particular SNP $k$ ; $t[\ ]$ : t-statistic measure of sexual dimorphism; Het: genome-wide mean heterozygosity. Population trait means or allele frequencies are indicated as such.

### What host traits affect parasite richness variation among individual hosts

Within each host population, we tested for correlations between individuals’ phenotypic traits and their parasite richness. Due to a shared environment and ancestry, any infection differences among fish within a population should stem from individual hosts’ phenotypic characteristics or genotype (i.e., Predictions 1-4). We focused exclusively on lake fish to exclude habitat differences among sites. We omitted watershed, which had no effect in M1. We used a Poisson General Linear Mixed Model (GLMM) to test whether per-fish parasite richness (*α_ij_*) depends on a random effect of population, standard length, sex, with population random effects of length and sex (i.e., trait*population interactions; Table 1, Model M2). For the subset of ∼30 fish with morphological trait data we ran a separate Poisson GLMM (M3) testing how *α_ij_* depends on population, sex, length, gill raker number and length, armor number, gape width, and all population random effects of these traits. For the subset of lakes with diet data (Table S1) we used a Poisson GLMM (model M4) testing for effects of population, sex, length, diet NMDS1 and NMDS2, prey richness, and all interactions with population (random slopes and intercepts). Full models for each of M2, M3, and M4, were evaluated against nested reduced models with AIC and likelihood ratio tests.

We carried out two analyses of the effects of host individuals’ genotype on individual parasite richness. First, we used genome-wide association (GWAS) to test each SNP’s effect on parasite richness (Model M5). Specifically, we used Poisson GLMMs to evaluate how individual parasite richness depends on population, sex, length, and focal SNP genotype. We included sex and length because they were substantial effects in the preceding model, and we do not wish to find loci contributing to variation in parasite richness via sex or size. We retained a population-specific random effect of sex, but not length, because only the former was supported in models M2-4. We applied M5 separately for each of 39,039 SNPs (restricting our attention to SNPs scored in at least 50 fish, with minor allele frequency exceeding 10%). We applied Holm corrections to the P-values.

Second, to test effects of whole-genome genetic diversity on parasite richness, we used Poisson GLMM (model M6) to relate parasite diversity to individual hosts’ genome-wide mean heterozygosity, with population as a random slope and intercept, a population random sex effect and a fixed effect of length. Due to small genotype sample sizes within populations, we omit random population effects of genotype or genome-wide heterozygosity.

### Does sexual dimorphism contribute to among-host variation

The effects of sex and the sex*population interaction on parasite richness were tested in models M2-M6. To evaluate possible phenotypic mechanisms of this dimorphism, we evaluated whether diet or ecomorphology dimorphism promotes sexually dimorphic infections. First, we calculated t-statistics for sex differences in parasite richness in each lake (t[*α_ij_*]) and t-statistics for sex differences in ecomorphology traits and diet (NMDS 1 and 2, and diet breadth [prey richness]). Then we used a linear model (M7) to test for a relationship between parasite richness dimorphism and dimorphism in size, morphology, and diet. We included only those traits that were, themselves, significantly sexually dimorphic and whose dimorphism varied significantly among populations. Although population constitutes the level of replication in this analysis, model M7 seeks to explain the magnitude of between-sex differences in infection (e.g., a source of within-population variation).

### What lake and fish traits drive parasite richness variation among host populations

We next tested for covariates of among-population variation in population-mean per-fish parasite richness (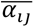). We focus on mean per-fish richness within populations, rather than aggregate parasite richness of a whole sample, because they are tightly correlated (r=0.58, P<0.0001; Figure S3), and the former is not biased by sample size and so does not require rarefaction. First, we tested for variation among habitats (lake, stream, estuary) while including a watershed random effect (model M8). Subsequently, we focus on lakes as the level of replication (omitting stream and marine sites). We tested the effect of geographic traits on 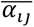 (Prediction 5), using lake area, ocean distance, and log lake size as fixed effects (model M9). Ocean distance is as-the-fish-swims distance along the river basin between the focal lake and the ocean. For the subset of lakes with ecomorphology data, we used an additional regression to test whether 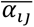 depends on population mean length, mean gape width, mean gill raker number, mean gill raker length, and mean armor plating (M10). For the subset of lakes that also have diet data, model M11 compared 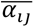 to mean diet (% benthic prey, prey richness, and NMDS1 and 2).

To test for genetic correlates of parasite richness (Prediction 3), we started with a population-scale GWAS analysis, using regression to relate 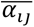 to the population allele frequency of a focal SNP (with watershed as a random effect, model M12). Again, we used Holm corrections for multiple comparisons. We regressed 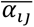 on population mean heterozygosity, 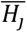 (M13).

Previous analyses of microsatellites suggest that heterozygosity in these lakes depends on lake characteristics (Caldera and Bolnick 2008), so we tested a final linear model relating mean heterozygosity to lake elevation, log area, and ocean distance (M14). Based on the results of models M9-14, we switched to a path analysis to account for causal relationships between predictor variables. We retained significant predictor variables from the linear models, keeping at least one variable per model M9-M14 (path diagram in Fig. 3).

## Results

### Among-host and among-population variation in parasite richness

Per-fish parasite richness (*αij*) varied from 0 to 10 species, with a mean of 2.28 species per host (sd=1.69, N=4375 fish; Fig. 1). Host population explained slightly less than half the variation (*η*^2^= 0.426; M1: host population effect, P<0.0001, details in Table S3), with no significant watershed effect. The remaining variation arose from among-individual differences in richness.

**Figure 1.**
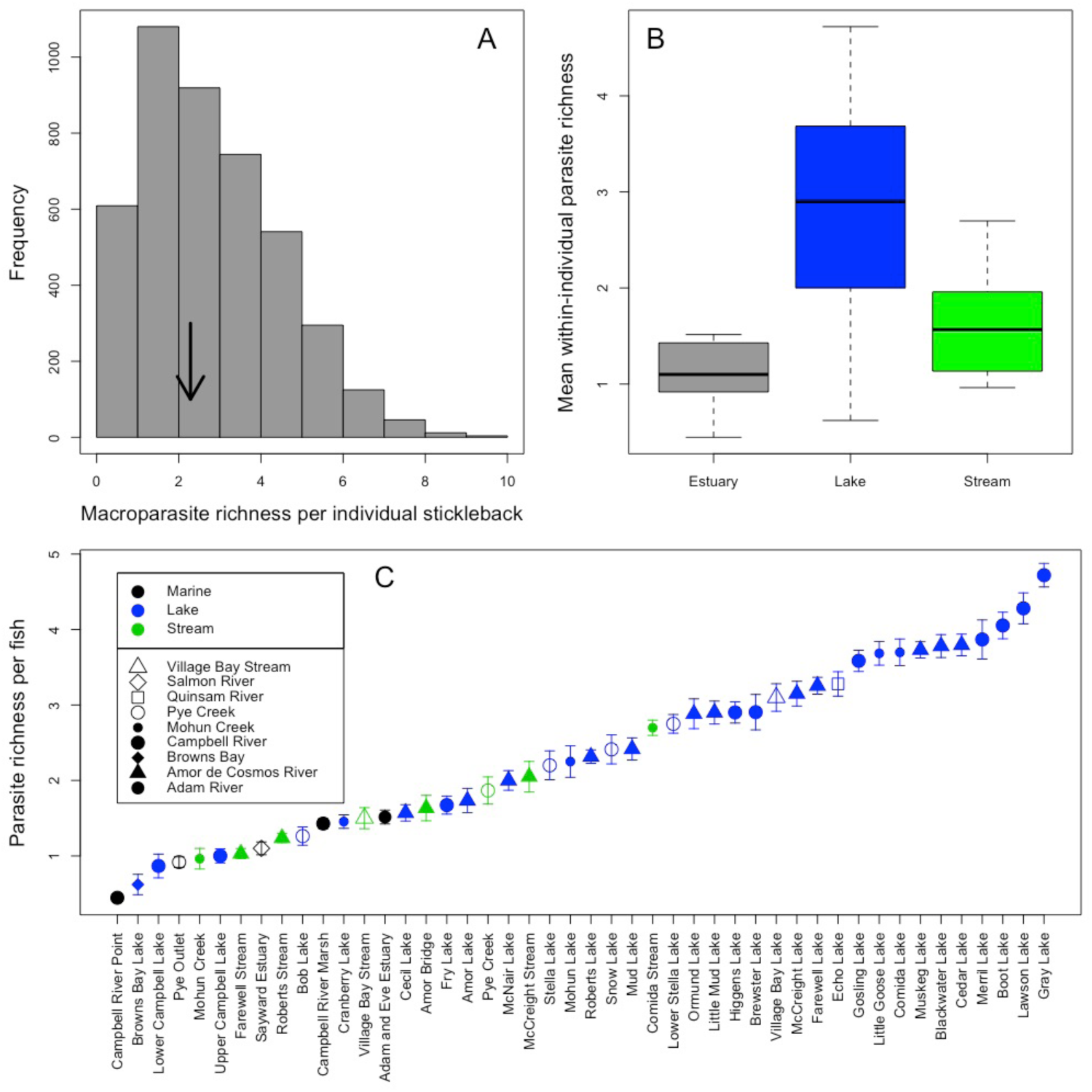
Per-fish parasite taxon richness (*αij*) varies substantially between (A) individual stickleback hosts, (B) habitats, and (C) populations. In (A) we use an arrow to indicate the distribution and mean per-fish parasite richness in the entire region, (B) shows the mean, 50% density, and 95% density of mean within-individual richness between habitats (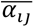), using host population as the level of replication. (C) plots 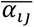 by population, sorted from least to most diverse. Points represent means with one standard error bars, color coded by habitat and symbols distinguishing different watersheds.

### Parasite richness covaries with individual ecomorphology

Individual hosts’ parasite richness was correlated with individual traits. AIC for M2 favored retaining fish length (standard normal; Z=11.02 P<0.0001), sex (Z=-1.22, P=0.22), a random effect of population, and a sex*population random slope. Larger fish tended to have higher parasite richness. Sex was not a significant main effect in M2, but there was a strong and significant sex*population interaction (P=0.0011), i.e., the degree and direction of sexually dimorphic parasite richness varies among populations (see *Sexual dimorphism in parasite richness* below). Models with a length*population interaction were not favored by AIC model selection (Fig S4).

At the individual host level, we found little evidence for ecomorphology or diet effects on parasite richness. Neither gill raker length, gill raker number, nor armor plate number correlated significantly with among-individual variation in parasite richness (M3; Table S3). Nor were there interactions between population and ecomorphology (LRT tests P>0.05). Neither host NMDS prey axis 1, axis 2, nor prey richness affected host-level parasite richness (M4; Table S3). There were no diet trait*population interactions. Only host size had significant effects.

Genomic analyses of individual parasite richness yielded no significant SNP-richness associations. Of 39,039 SNPs tested with model M5, 1265 were significant at P < 0.05 and 21 at P < 0.001, but no more than false discovery rate expectations. We found no effect of individual genome-wide heterozygosity on parasite richness (M6, Poisson GLMM; Z=-1.36, P=0.174).

### Sexual dimorphism in parasite richness

Parasite richness was significantly dimorphic in 17 of the 42 populations (Fig. 2A). Of these 17 dimorphic populations, females had higher richness than males in 13 sites (M2, sex*site interaction, χ^2^=21.68, df=2, P<0.0001). The magnitude and direction of sex dimorphism in parasite infection was predictable using ecologically relevant traits (Fig. 2B,C; M7). Parasite richness was higher in whichever sex had larger body size (t=3.759, P=0.0009) and whichever sex had lower NDMS2 scores (t=-2.48, P=0.021). These lower NDMS2 scores indicate greater chironomid larval consumption but fewer large prey like *Gammarus*, ceratopogonid larvae, and stickleback eggs (Table S3). NDMS1, which negatively covaries with the proportion of benthic prey (r=-0.51, t=-16.8, P<0.0001), was significantly dimorphic but did not predict dimorphism in parasite richness (t=-1.2, P=0.25).

**Figure 2.**
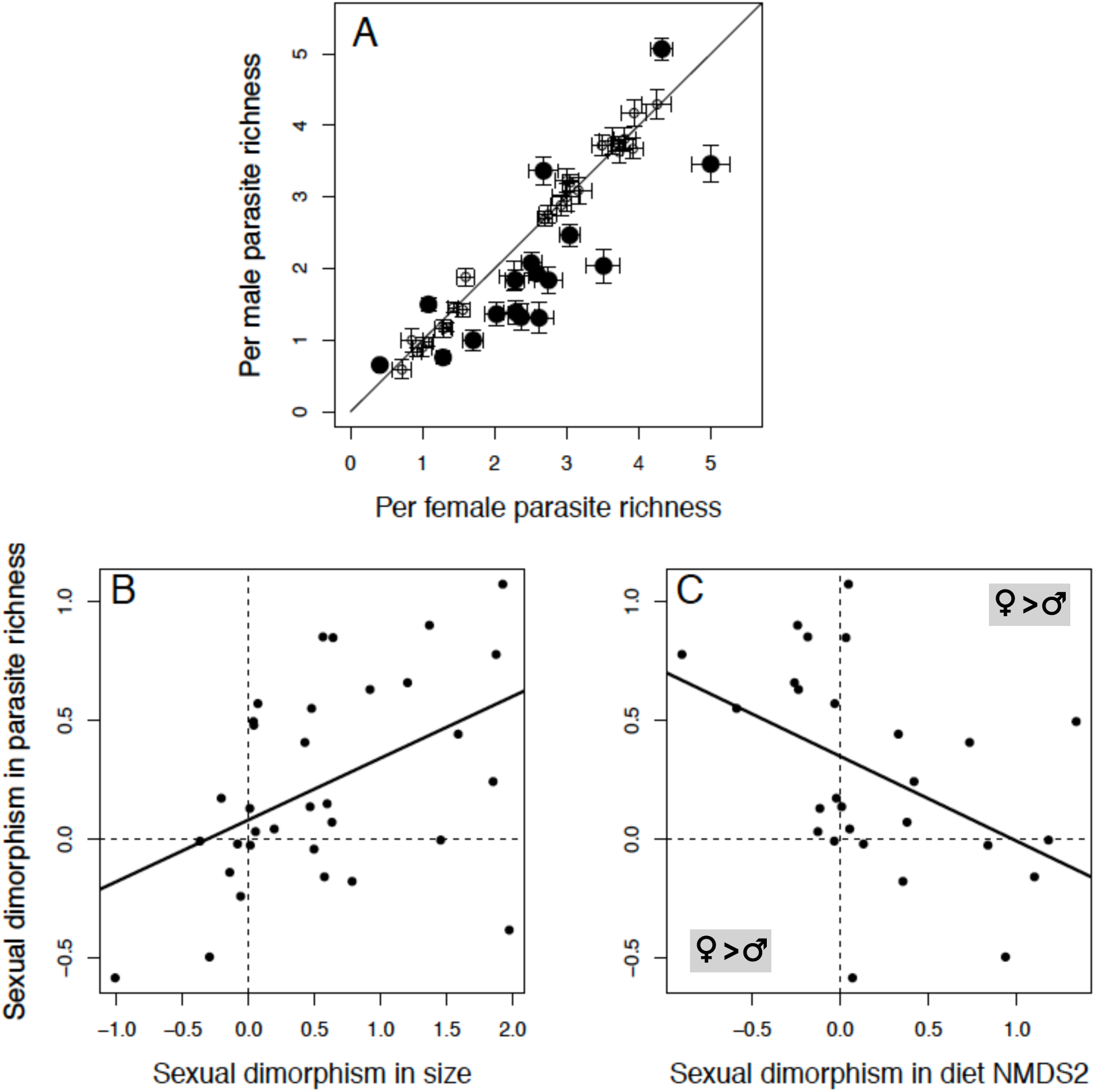
A) Sexual dimorphism in parasite richness is observed in 17 of the 41 sites for which we had at least 10 fish per sex. Large solid points indicate significantly dimorphic populations that deviate from the 1:1 diagonal line. Smaller open circles are not significantly dimorphic populations. Of the 17 dimorphic cases, in 13 populations females carried more diverse infections whereas in 4 the males had more diverse infections. This variation in dimorphism direction and magnitude is associated with the direction and magnitude of dimorphism in B) host size, and C) host diet. In each panel, dimorphism is calculated as female mean minus the male mean. So, positive values denote populations in which females were larger, ate prey scoring high on diet NMDS2 (more ceratopogonids, gammarus, and stickleback eggs but fewer chironomids), and had higher parasite richness. The trends in B and C are also observed if we use Shannon-Weaver diversity of parasites (P = 0.0075 and 0.0022, respectively).

### Among-population differences in parasite richness

Mean per-fish parasite richness (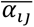) spanned an order of magnitude across fish populations (Fig. 1C; model M1 P <0.001)— 0.44 ±0.05s.e. taxa per fish in marine fish captured at Campbell River Point, to 4.72 ±0.16s.e. in Gray Lake. Richness varied by habitat (M8, habitat effect χ^2^=19.0, df=2, P<0.0001), being on average 2.8-fold higher in lake stickleback than in anadromous fish (Fig. 1B; χ^2^=14.29, df=1, P=0.0002) and 1.8-fold higher than stream fish (χ^2^=7.79, df=1, P=0.0052). Stream stickleback had, on average, 1.5 times as many parasite taxa as marine fish (χ^2^=3.63, df=1, P=0.056). Watershed had no detectable effect, so we do not consider it further.

Lake geography partly influenced among-lake variation in mean parasite richness (M9). We found no support for effects of elevation (t=-0.58, P=0.56) or lake depth (t=0.6, P=0.55), weak support for an effect of log surface area (t=-1.84, P=0.076), and a significant positive effect of ocean distance (t=2.56, P=0.016; Fig. S5). A few post-hoc observations are worth noting. First, although lake depth had no linear effect on parasite richness, visual inspection of the data suggested a quadratic relationship (depth t=4.18, P=0.0001, depth^2^ t=-3.7, P=0.0006, Fig. S6). Log lake surface area likewise had a quadratic effect on mean richness (area t=2.11, P=0.043; area^2^ t=-2.64, P=0.013, Figure S7). Last, although elevation had no main effect in M9, inspection of the data suggested an interaction with lake area, which was confirmed with a post-hoc linear model (elevation t=3.82, P=0.0006, log surface area t=3.00 P=0.0055; elevation*area t=-3.46, P=0.0016, Fig. S8).

We found support for some host-population trait means explaining among-lake differences in parasite richness (M10, fish length t=2.8, P=0.008; size-adjusted gape width t=2.06, P=0.048; gill raker number t=1.87, P=0.072; size-adjusted gill raker length t=-0.026, P=0.979). AIC supported a simpler model in which mean parasite richness () increased with mean fish length and gape width, but decreased with mean gill raker number (all typical benthic ecomorph traits). More benthic-feeding populations (higher NMDS1) tended to have more parasites per fish but this was marginally significant (P=0.07, M11). NDMS2 and mean prey richness had no effect (P=0.42 and 0.41 respectively).

Genome-wide association mapping (M12) revealed no significant (after Holm correction) SNP effects on host-population parasite richness. Mean genome-wide heterozygosity had no detectable association with population mean parasite richness (M13, t=0.925, P=0.364).

The variables considered above are likely to be inter-related. For example, heterozygosity increases with lake area (Caldera and Bolnick 2008) but decreases with distance from the ocean and elevation (M14; P=0.029, 0.012, <0.001 respectively). To account for this covariance among predictors, we used path analysis to partition direct and indirect effects (Figs. 3&4, Table S4), taking the significant effects from the 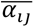-based linear regressions above. The path analysis explained 45.6% of the variation in mean per-fish parasite richness among lakes. Lake area had no significant direct effect on parasite richness (r=0.053, P=0.822), but had indirect effects via fish heterozygosity and diet. Specifically, larger lakes have more genetically diverse fish (r=0.366, P=0.011), and heterozygosity has a positive effect on parasite richness (r=0.360, P=0.038). Stickleback in larger lakes also consumed a smaller proportion of benthic prey (r=-0.777, P<0.001), which in turn conferred higher parasite richness (r=0.570, P=0.013). The indirect negative effect of lake size via fish diet exceeded its positive effect via fish heterozygosity (r=-0.311 versus 0.130). In simple bivariate correlation tests, lake distance from ocean had a positive effect on parasite richness (r=0.310, P=0.017). However, this positive correlation is mediated via an indirect positive effect of ocean distance on heterozygosity (Table S4; r=0.553, P=0.002), which has a positive partial correlation with parasite richness. Lakes farther from the ocean also had larger fish (r=0.52, P=0.001), but in this analysis, mean per-lake fish size had no further relevance to fish diet (r=0.16, P=0.111) or parasite richness (r=0. 075, P=0.635). Last, higher elevation lakes had less heterozygous fish (r=-0.74, P<0.001), indirectly reducing richness.

**Figure 3.**
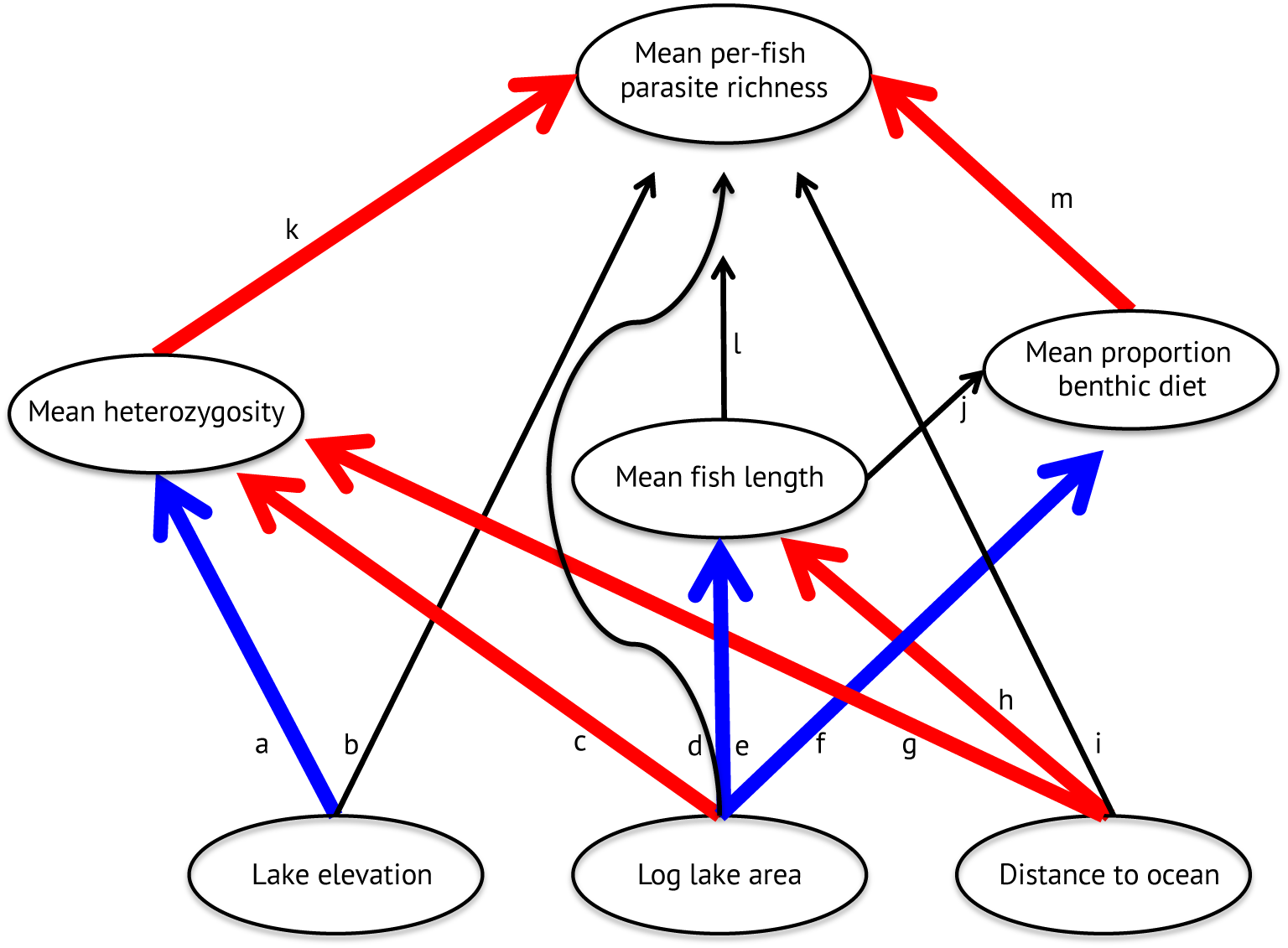
Diagram of the path analysis of among-lake variation in mean parasite richness. Thin black lines were non-significant effects. Thick red and blue lines, respectively, indicate significant positive and negative effects. Partial correlations are labeled by letters corresponding to the effects plotted in Fig. 4.

**Figure 4.**
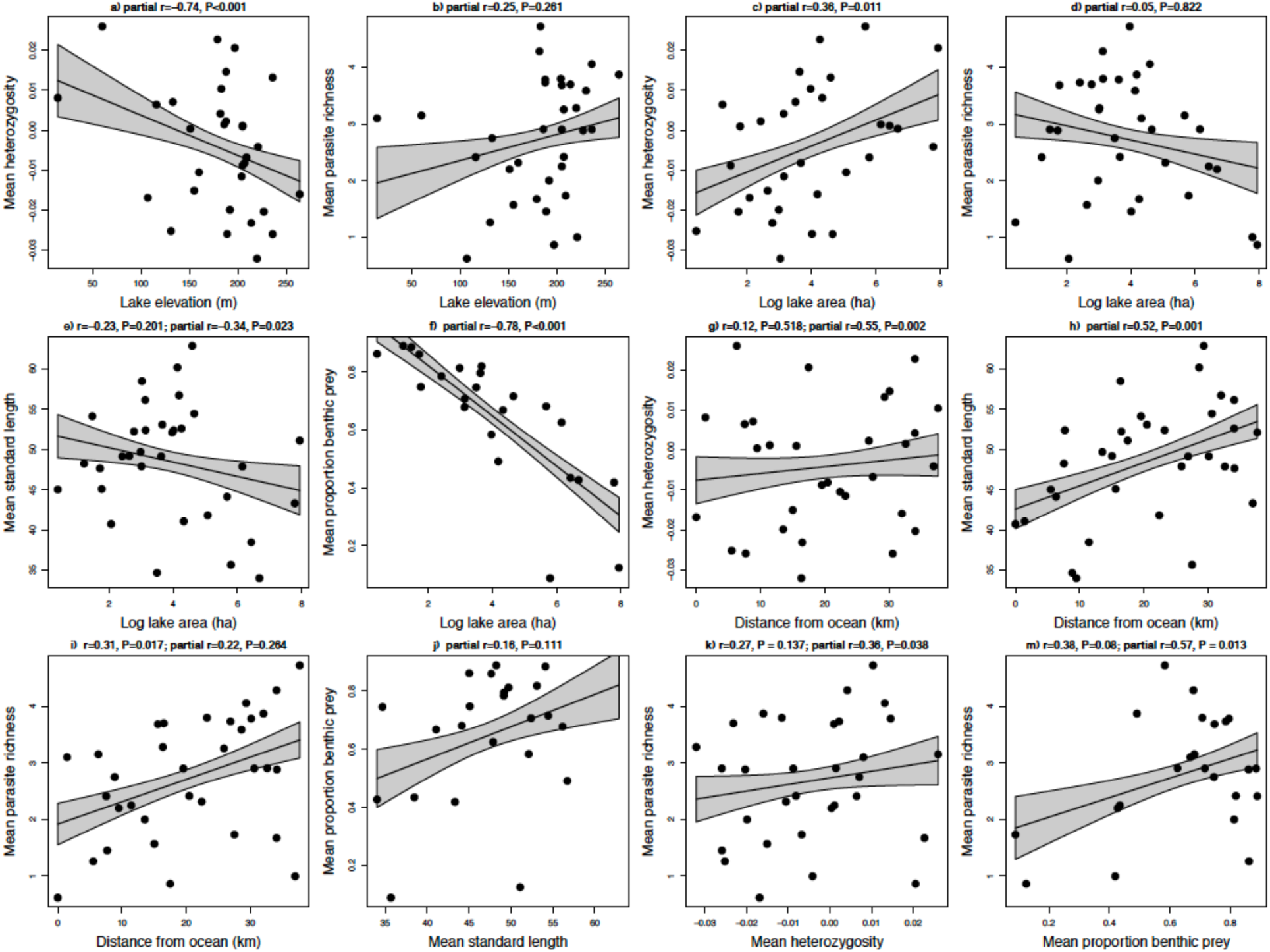
Scatterplots of the correlations tested in the path analysis illustrated in Fig. 3, with linear regression fit lines and confidence intervals for the raw data. Each panel is labeled (a) through (m) to match the path arrows in Fig. 3, omitting path l which was not significant. Above each panel we provide the partial correlation and its significance, from the path analysis (Table S4 for details). In cases where the simple bivariate correlation yields a different trend than the path analysis (e.g., when one is significant and the other is not), we provide both the bivariate correlation then the path results.

## Discussion

Parasites form diverse multi-species assemblages in nested metacommunities across individual hosts, and among host populations (Seabloom *et al*. 2015). Key questions in parasite ecology, and ecology more broadly, are (1) what processes dictate this metacommunity diversity, and 2) are these processes scale-dependent? Here, we present a case study using macroparasites from a stickleback metapopulation to show that parasite richness varies widely among host individuals and across host populations. In both cases, the variation in richness spans an order of magnitude. We have identified ecological factors contributing to this variation in parasite diversity and have shown that mostly different factors act at small and large spatial scales (summarized in Fig. 5).

**Figure 5.**
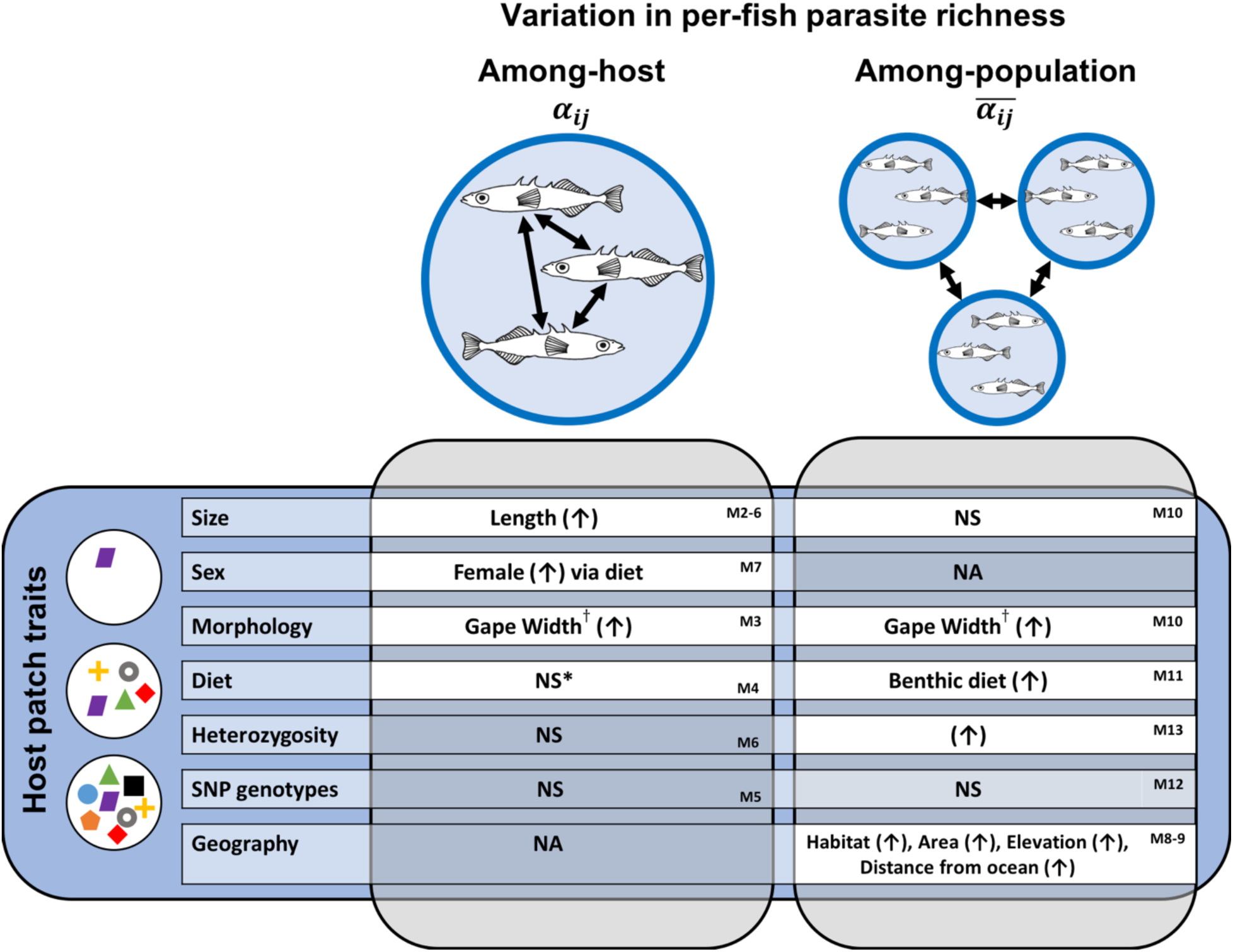
Summary of inferences at the among-host and among-population scales, indicating statistically significant effects (with check-marks), non-significant effects (NS) and effects not operating at a given scale (NA). * Sex differences in diet contribute to between-sex differences in parasite diversity within populations

### Individual scale: Why does parasite richness differ among host individuals?

Individual hosts within a population experience approximately similar abiotic conditions and biotic prey communities, at least for geographically well-defined populations like fish within a lake. Thus, variation in parasite richness among host individuals is either stochastic or due to differences in individual traits. We confirmed that stickleback individual size, gape width, diet, and sex were correlated with individual parasite richness, in sometimes subtle ways, but did not find any effect of genotype or heterozygosity.

Larger individual stickleback carried more parasite taxa, as in other fish species (Calhoun *et al* 2018). Larger fish may be older, having more time to accrue infections or having senescing immune systems (Zelmer 2014). Or, larger stickleback might also be capable of eating more prey due to larger stomachs, increasing overall intake rates of parasites. A third possibility is that larger fish eat particular kinds of prey that promoting parasite diversity. For instance, in stickleback and many other fish, larger individuals are more likely to be benthivorous. Although we do not have an a priori expectation that benthic-feeding fish would have greater parasite richness, we do also find that fish with larger gapes (a typical benthic trait) also have higher parasite diversity. Moreover, our sexual dimorphism analysis suggests that parasite richness is higher in whichever sex has the more benthic diet and is larger. Thus, we conclude that individual size and trophic ecology affect individual hosts’ parasite richness. This confirms that habitat patch characteristics (i.e., individual host traits) modify community assembly, likely by modulating the rate of colonization.

Diet effects are consistent with previous studies in other species that found that individual diet influences parasite infection (Amundsen *et al*. 2003; Stutz *et al*. 2014; Cirtwill *et al*. 2015; Hayward *et al*. 2017), and with the general notion that parasite communities are structured by colonization dynamics (Worthen and Rohde 1996; Zelmer 2014).

Males and females differ in diet and microhabitat use in most species (Shine 1991; McGee and Wainwright 2013; Reimchen *et al*. 2016). Thus, individual encounter rates with different parasites should vary by sex (Reimchen and Nosil 2001). In stickleback, we found that sex-biased parasite richness covaries with sexual dimorphism in ecomorphology traits, implying a role for encounter filters. Across all lakes, females tended to be larger than males, consume more large prey, and have higher parasite richness. In lakes with larger dimorphism in body size or diet, parasite richness was also more dimorphic. The exceptions support our conclusions: in a few lakes where diet or size dimorphism is reversed, parasite richness dimorphism is also reversed. Sexual dimorphism in immune function (Zuk 1996; Nunn *et al*. 2009), which we did not measure, could contribute to the residual variation in sex dimorphism of parasite richness.

We found no single-locus or genome-wide genetic control of per-fish parasite richness (*αij*). This is perhaps because parasite richness is an aggregate measure arising from many parasite species’ interactions with the host, which might be controlled by separate gene(s). In the companion study (Bolnick et al., Manuscript), we report many genetic associations with infection by each common parasite. But, because these associations tend not to overlap, there is no overall genetic effect on parasite richness.

### Population scale: Why does parasite richness differ among host populations?

Parasite richness varied substantially among host stickleback populations with habitat, geography, ecomorphology, and genetic diversity. The strongest effect was habitat: parasite richness was nearly three times higher for lake stickleback than their anadromous relatives; stream fish were intermediate, consistent with previous studies (e.g. Eizaguirre *et al*. 2011; 2012). Given that stickleback are relatively recent (post-glacial) colonists of freshwater, this higher infection rate indicates that the colonization process increased infection risks, rather than providing a means of escape from historical enemies (Grunberg *et al*. 2019). It remains unclear to what extent these habitat differences are a function of abiotic conditions, availability of intermediate or terminal hosts, or habitat differences in dilution effects (e.g., higher non-host fish species richness in the ocean; Becker *et al*. 2014).

For lake populations, lake geography affected parasite richness, which was higher in mid-sized lakes (with intermediate depths), farther from the ocean, and at higher elevation. Some geographic effects acted indirectly via host population traits. For example, stickleback in larger lakes have more limnetic diets on average (Lavin and McPhail 1986), and this diet shift is associated with reduced parasite richness; i.e., lake size had a net negative indirect effect on parasite richness via diet.

This negative effect was lessened somewhat by a positive but weaker indirect effect of lake size through host heterozygosity. Larger lakes support more genetically diverse stickleback populations (Caldera and Bolnick 2008, confirmed in this study). This increase in mean heterozygosity was associated with higher mean per-fish parasite richness, inconsistent with the oft-cited immunological benefit of genetic diversity (Joly *et al*. 2008; Kaunisto *et al*. 2013; Poulin *et al*. 2000). This surprising effect might be explained if we consider a reversed cause-effect relationship: richer parasite communities might select for greater genetic diversity. But, such selection is unlikely to affect genome-wide heterozygosity, because (1) most SNPs should be effectively neutral, and (2) heterozygosity is mostly a function of lake geography.

Heterozygosity was also influenced by lake elevation and distance to ocean. Higher-elevation lakes tend to have lower heterozygosity, consistent with expectations of a stronger bottleneck during colonization. Path analysis suggests that elevation reduces parasite richness indirectly via reduced heterozygosity, but there was no direct elevation-richness correlation. Likewise, lake distance from the ocean only had an indirect effect on richness via heterozygosity.

Our results do not conform to general predictions from island biogeography and basic metacommunity theory. All else equal, communities should be more diverse on habitat patches that are larger, or closer to other such patches (MacArthur and Wilson 1963), as demonstrated for human parasites on islands in Jean *et al*. (2016). Larger lakes are akin to larger islands, but in our study, lake size had no net relationship to parasite richness due to conflicting effects via host diet and heterozygosity (noted above). Furthermore, we find that more isolated stickleback populations with longer distances to the ocean have richer, rather than poorer, parasite communities. We hypothesize that these unexpected effect directions could be driven by host evolutionary genetics.

### Scale-dependent and -independent factors affecting parasite richness

Thus far, we have discussed each spatial scale separately. To what extent do the variables structuring parasite diversity generalize across scales (Fig. 5)? Only two traits showed similar effects across scales. Larger individual fish had richer parasite infections, and populations with larger average fish had higher mean parasite richness. We also detected a marginally significant positive effect of gape width on parasite richness at both scales, which suggests the trend at each scale is real rather than Type I error. These effects of size and gape confirm Prediction 1.

Direct measures of diet had inconsistent effects across spatial scales. At the larger spatial scale, stickleback populations that were (on average) more benthivorous (lower NDMS1) had richer average per-fish parasite communities. No such trend was found at the individual scale. Diet sexual dimorphism (acting within populations) did correlate with parasite richness dimorphism, but this involved NDMS2. Thus, diet does influence parasite richness, but does so inconsistently across scales. Prediction 2 is supported only at the among-population scale.

A scale-specific relationship was also seen between genomic heterozygosity and parasite richness. Some studies suggest that heterozygosity might increase a host’s repertoire of immunological tools, thereby lowering richness (Coltman *et al*. 1999; Arkush *et al*. 2002; Whitehorn *et al*. 2011). Indeed, fish species with higher heterozygosity had fewer parasite species (Poulin *et al*. 2000; Joly *et al*. 2008; Kaunisto *et al*. 2013). Our data did not support such a trend at the individual-level, despite high statistical power, partly refuting Prediction 3. The lack of an individual-level effect undermines the notion that heterozygosity acts through individual immunogenetic diversity. We did, however, find a population-level effect of heterozygosity, though in an unexpected direction (parasite richness increased with host population mean heterozygosity).

So, why does heterozygosity, an individual trait, matter at the population scale? One possibility is that larger host populations can both support more diverse parasite communities and maintain higher heterozygosity. Parasite diversity can be related to host population size, following a species-area relationship (Bagge *et al*. 2004; Zelmer 2014). However, our path analysis found no support for a direct effect of lake size on parasite richness. An alternative explanation is that fish species exposed to more diverse parasites might be subject to stronger balancing selection and evolve higher heterozygosity (Hamilton 1982; Poulin *et al*. 2000; Berenos *et al*. 2010). But, this adaptationist interpretation should not impact genome-wide heterozygosity, only those loci involved in immune defense and sites linked to them.

### Conclusions

Because parasites are such a large component of biological diversity and have large effects on the communities in which they are embedded (Lafferty *et al*. 2008), the field of ecology needs to understand the biological processes regulating the distribution, abundance, and diversity of parasites. Here, we present a case study illustrating the highly multivariate and scale-dependent nature of these processes. The macroparasite community of threespine stickleback is structured by among-individual variation in host sex, size, morphology and diet, and among-population differences in host size, morphology, diet, genetic diversity, and habitat. Some of these variables (size, gape width) act consistently across individual- and population-scales. In general, it seems that more benthivorous stickleback have richer parasite communities, a trend observed between sexes, and among populations. Other variables are scale-dependent and contribute to parasite differences only among individual hosts (sex) or only among host populations (heterozygosity, habitat). Such scale-dependent effects are expected in metacommunities of all kinds (Leibold *et al*. 2004). This scale-dependence may help explain inconsistent results among studies conducted at disparate scales. The implication of these findings is that studies of species distributions and abundances need to draw scale-specific inferences.

## Acknowledgements

We thank C. Harrison, T. Rodbumrung, and T. Ingram, for assistance with field work. J. Day assisted with parasite data collection. R. Grunberg commented on the manuscript. The research was supported by the Howard Hughes Medical Institute through an Early Career Scientist position to DIB, NSF (DEB-1144773) to DIB, and a NSF Graduate Research Fellowship to EJR.

---

**Supplementary Figure S1.**
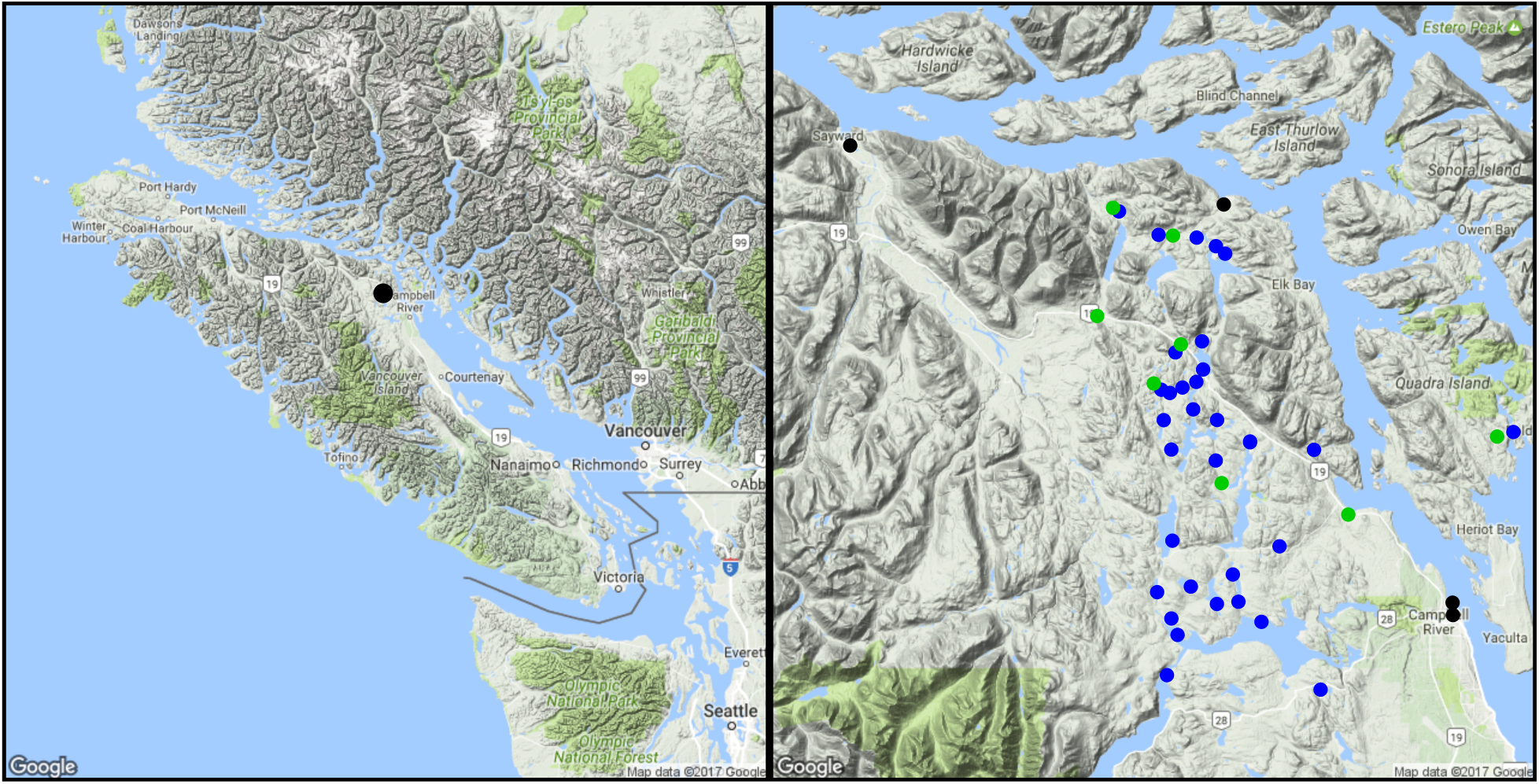
A) Map of Vancouver Island, British Columbia, with a dark point showing the location of the inset map B) on which we plot black, green, and blue dots to indicate marine, stream, and lake populations sampled. The 5^th^ marine site is farther northwest than shown in (B)

**Supplementary Figure S2.**
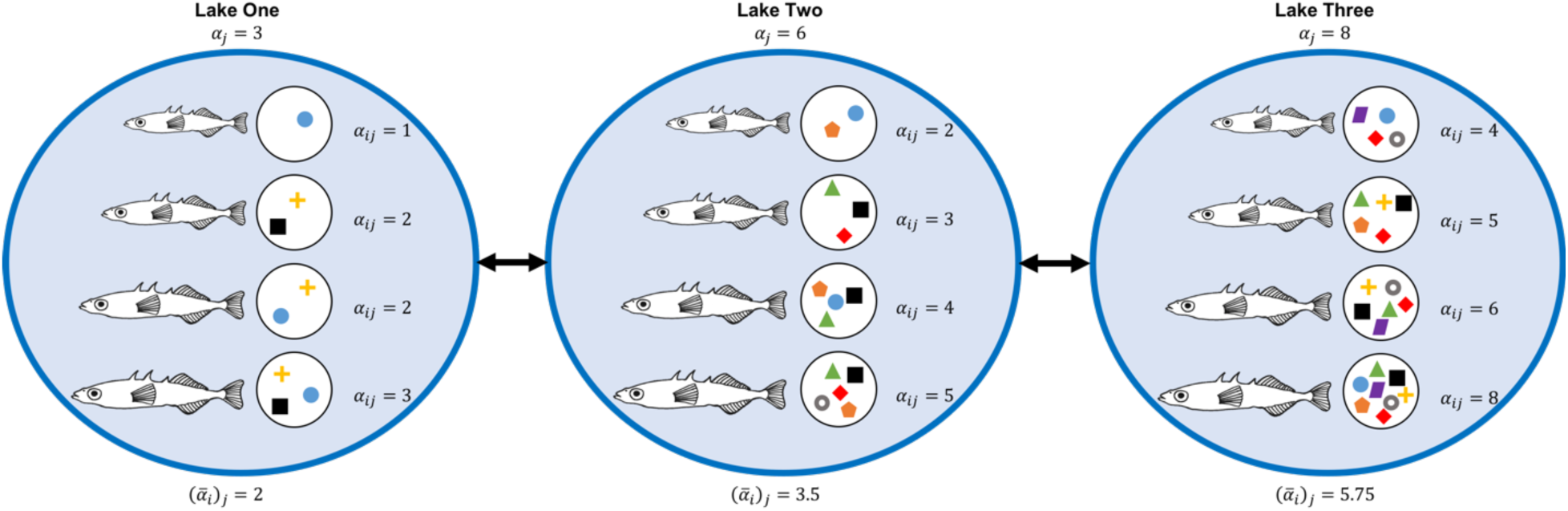
Illustration of variation in parasite richness among individual fish and among populations. Each blue circle represents a lake containing phenotypically varying hosts (arranged from small to large). Each host carries parasites (indicated by symbols in small circles. Individuals differ in parasite richness (*αij*). The populations differ in mean parasite richness 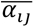, the variance in richness among individuals (*σ*(*α_i_*)_*j*_), and the total parasite richness per lake (*α*\*_j_).

**Supplementary Figure S3.**
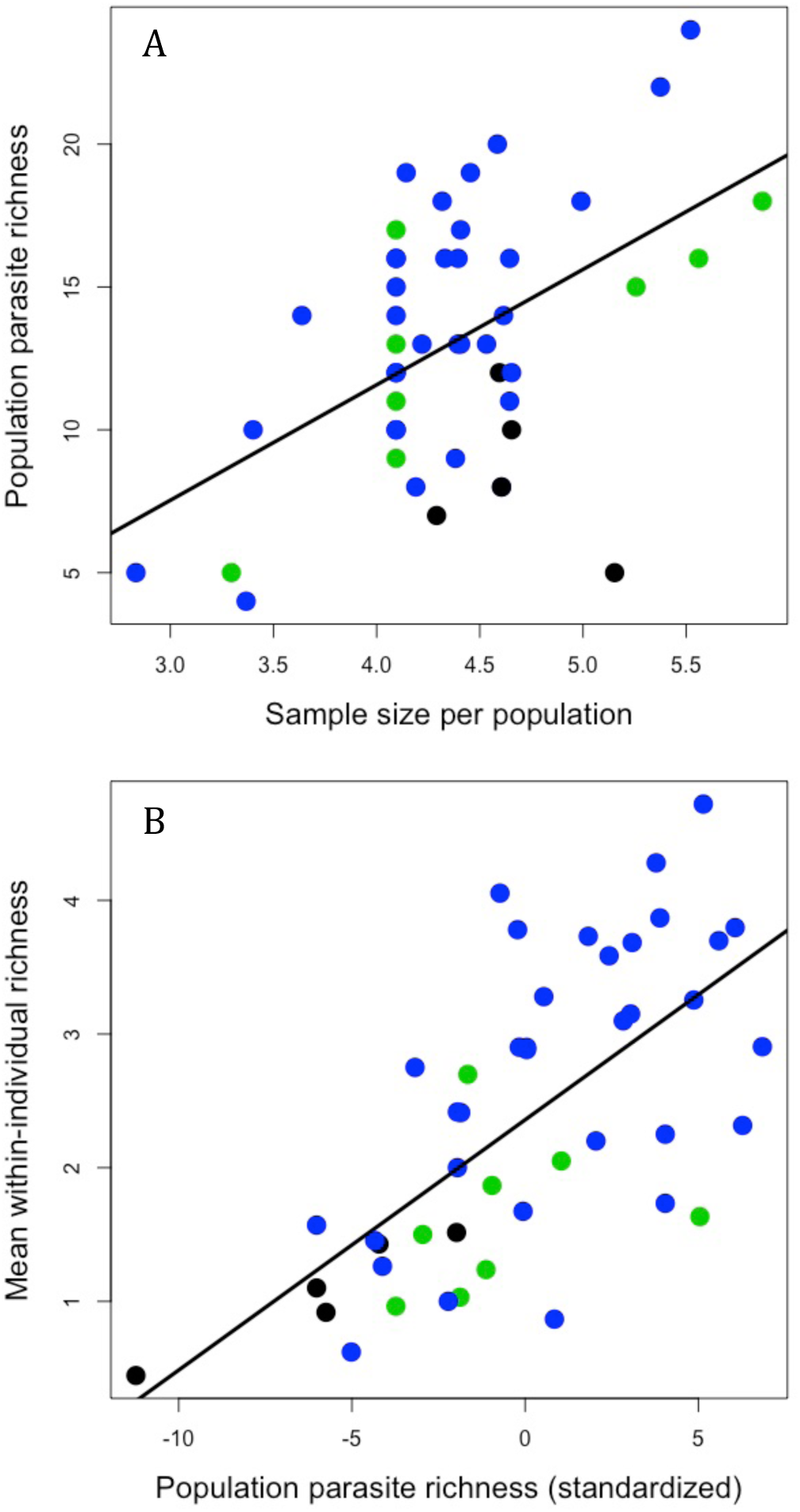
A) Population-wide parasite richness (*α*\*_j_) increased with the number of sampled fish (shown on a log scale), unlike mean per-fish richness. B) The two measures of parasite richness, (*α*\*_j_ and 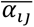) are highly correlated. Blue, black, and green points represent lake, marine, and stream sample locations. Here, we use richness standardized by individual host mass.

**Supplementary Figure S4.**
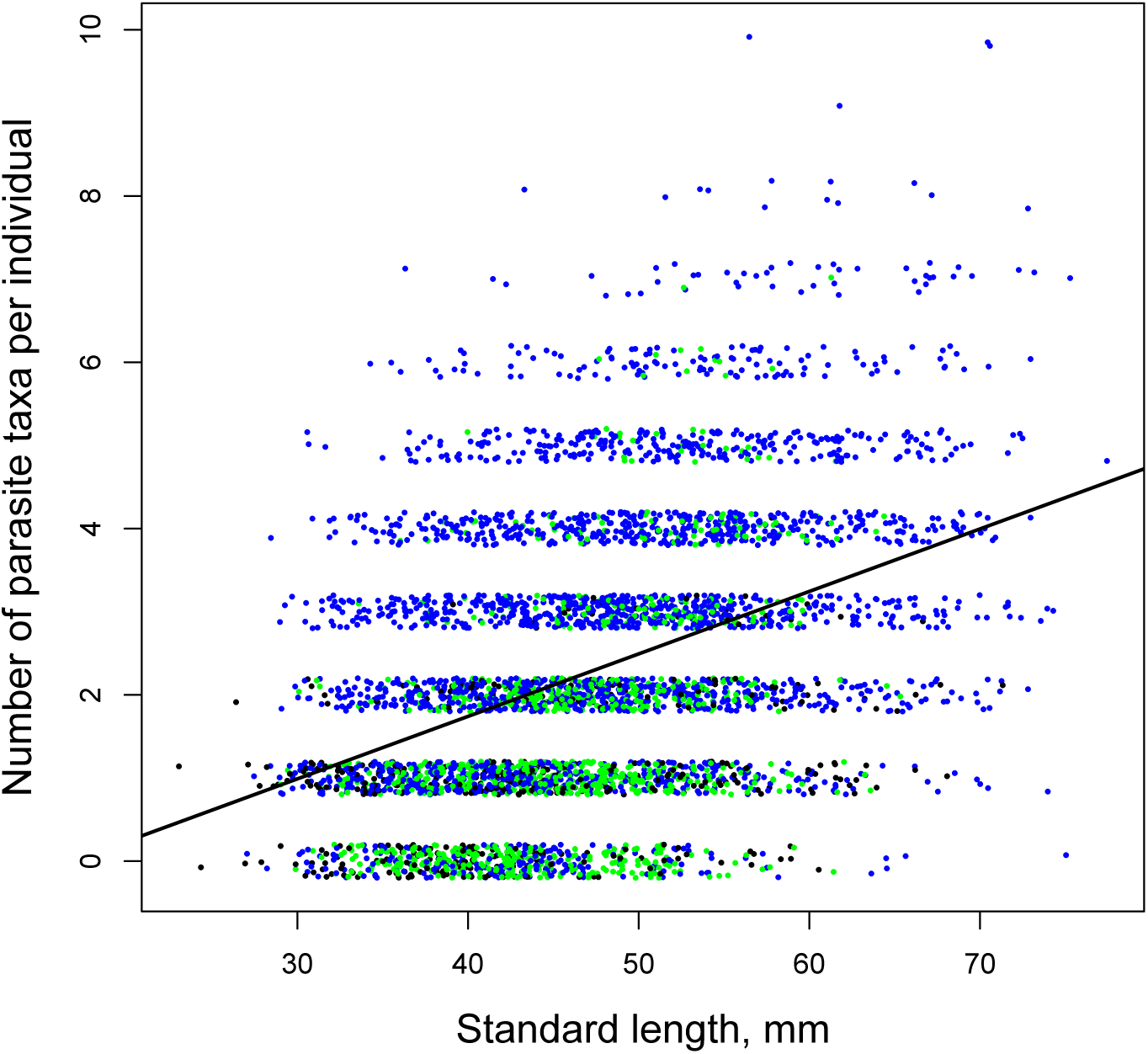
Parasite richness as a function of host length. Points are individual fish. Blue, green, and black dots correspond to lake, stream, and marine fish. Some jitter is added to the y axis to distinguish points.

**Supplementary Figure S5.**
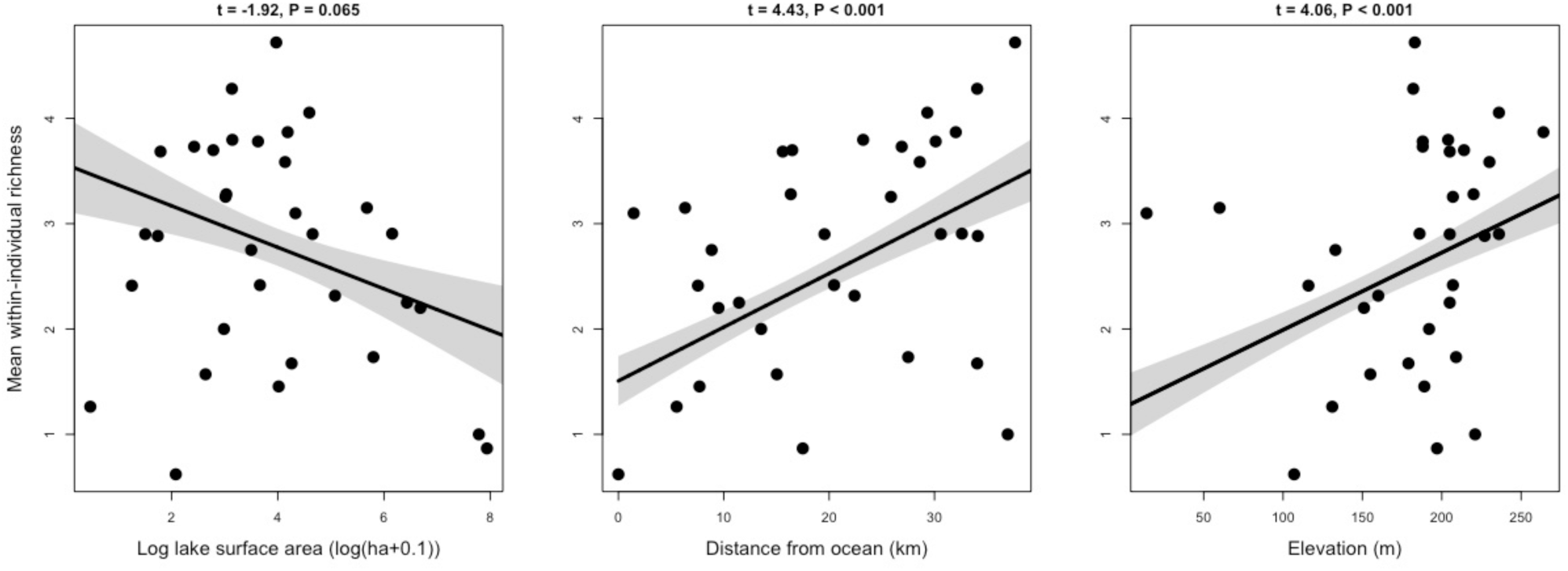
Effects of lake characteristics on population mean parasite richness per fish, using only lakes as the level of replication.

**Supplementary Figure S6.**
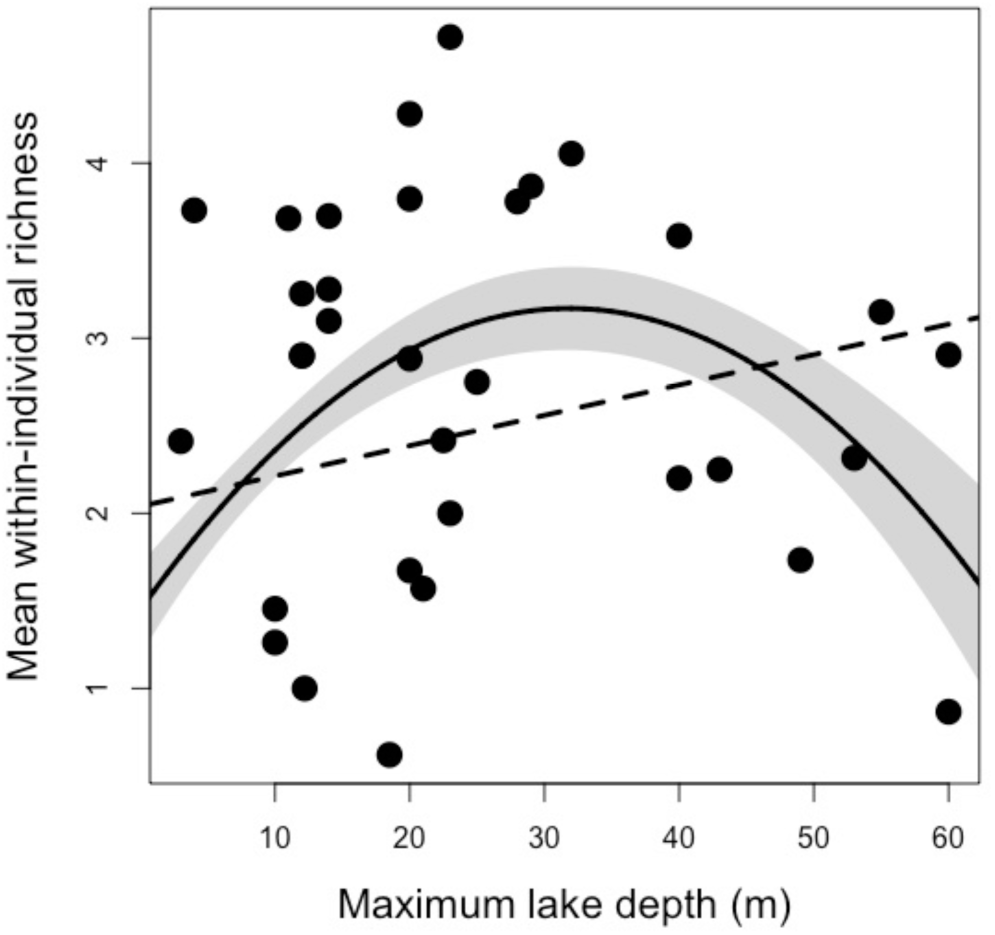
Quadratic (solid line, with 1 standard error confidence interval shaded) and linear (dashed line) regression estimates of the effects of maximum lake depth on population mean per-fish parasite richness. The quadratic effect is strongly significant whereas the linear is not, but was only tested after visual inspection of the data, so is a post-hoc analysis.

**Supplementary Figure S7.**
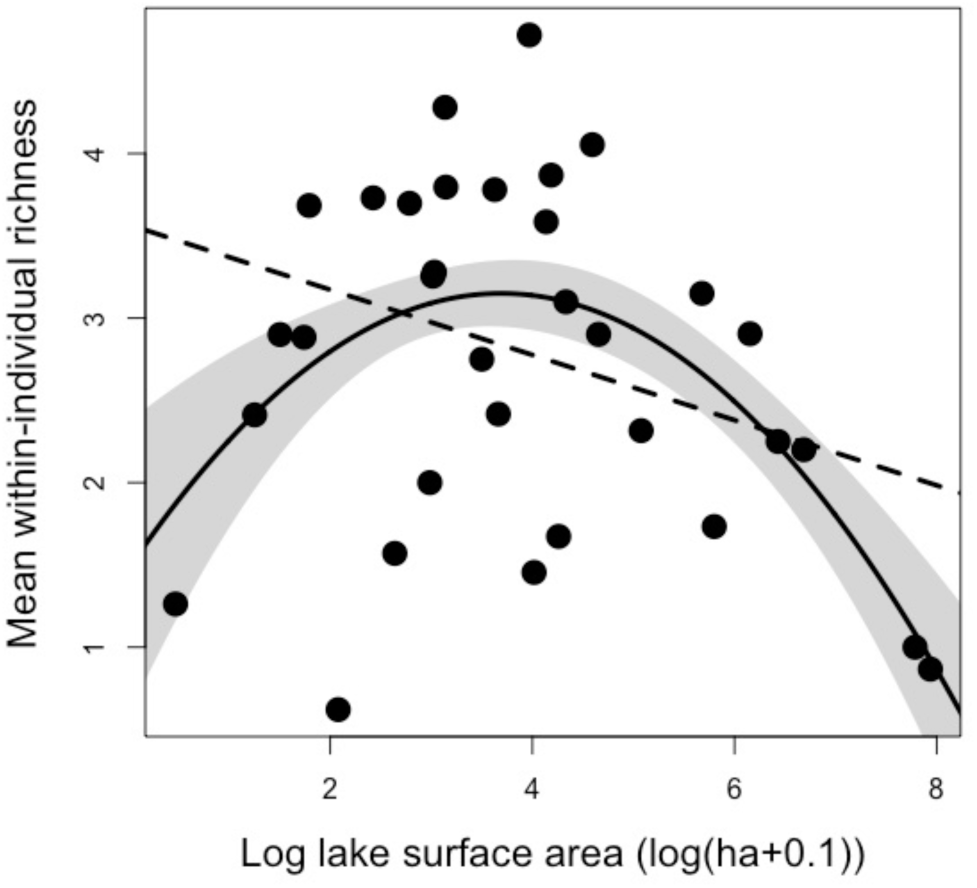
Quadratic (solid line, with 1 standard error confidence interval shaded) and linear (dashed line) regression estimates of the effects of log lake area on population mean per-fish parasite richness. The quadratic effect is significant whereas the linear is not (Figure S5), but was only tested after visual inspection of the data, so is a post-hoc analysis.

**Supplementary Figure S8.**
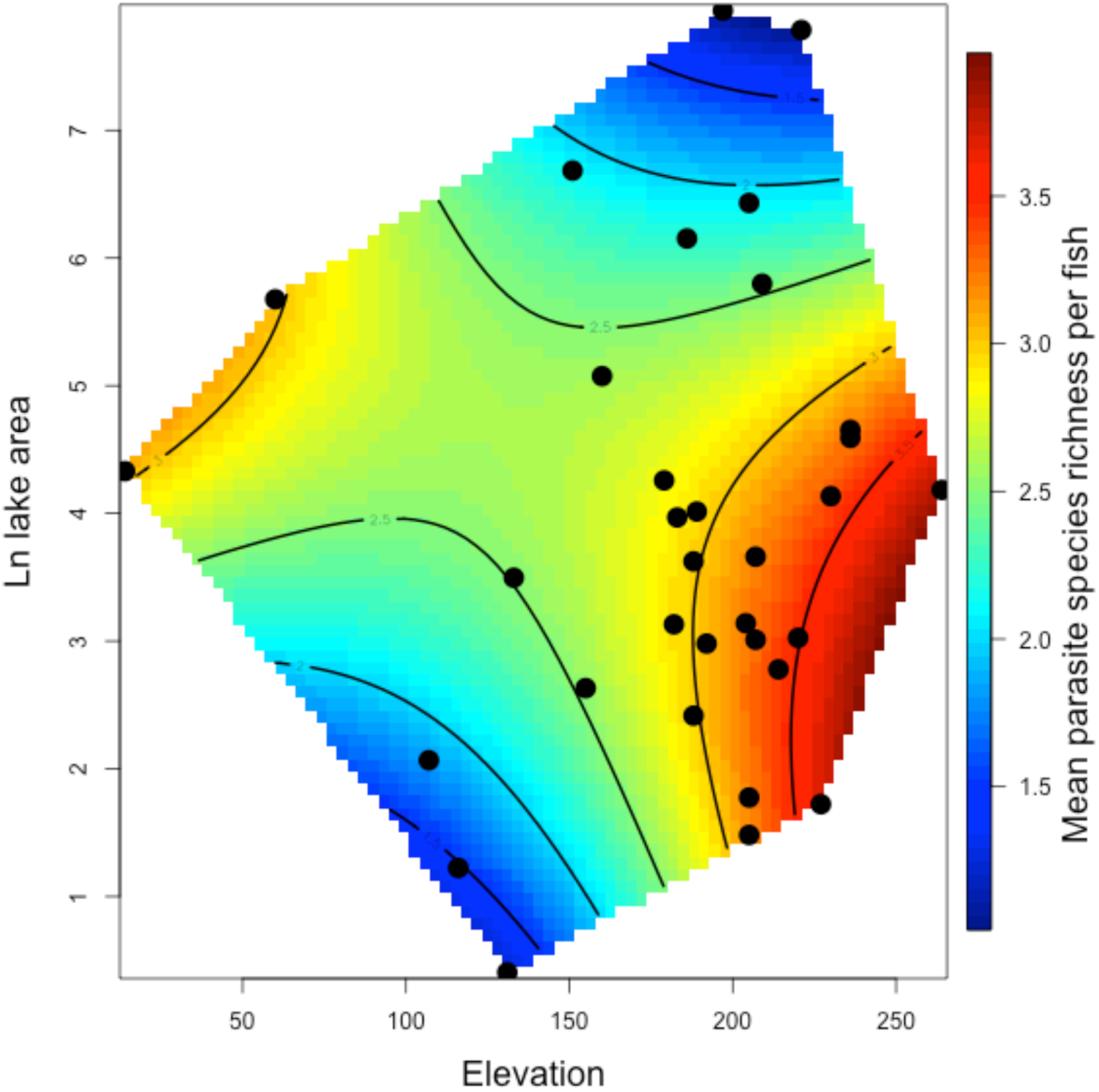
An interaction effect between elevation and log lake area on mean per-fish parasite richness: for low elevation lakes (e.g., <150 m), lake area increased parasite richness, whereas for high-elevation lakes the reverse was true. This interaction was identified and statistically tested after visualizing this trend, so is a post-hoc analysis.

**Table S1.** Summary of sampled populations, locations, sample sizes, and mean parasite taxon richness.

|  |  |  |  |  |  |  |  |  | Mean |  |
| --- | --- | --- | --- | --- | --- | --- | --- | --- | --- | --- |
|  |  |  | Sample | Trophic |  | ddRAD |  |  | parasite | SD parasite |
| Site name | Longitude | Latitude | size | morphol. N | Diet N | N | Habitat | Watershed | richness | richness |
| Adam and Eve |  |  |  |  |  |  |  |  |  |  |
| Estuary | -131.9426 | 50.4012 | 99 | 99 | 0 | 10 | Estuary | Adam River | 1.52 | 0.91 |
| Amor Bridge | -125.6639 | 50.2543 | 60 | 30 | 23 | 10 | Stream | Amor River | 1.63 | 1.30 |
| Amor Lake | -125.5791 | 50.1581 | 60 | 30 | 30 | 10 | Lake | Amor River | 1.73 | 1.25 |
| Blackwater Lake | -125.5878 | 50.1794 | 82 | 30 | 25 | 16 | Lake | Amor River | 3.78 | 1.38 |
| Bob Lake | -125.5281 | 50.3045 | 80 | 61 | 61 | 10 | Lake | Pye River | 1.26 | 1.09 |
| Boot Lake | -125.5270 | 50.0467 | 93 | 93 | 0 | 0 | Lake | Campbell River | 4.05 | 1.70 |
| Brewster Lake | -125.5779 | 50.0924 | 63 | 30 | 28 | 10 | Lake | Campbell River | 2.90 | 1.86 |
| Browns Bay Lake | -125.4159 | 50.1580 | 29 | 0 | 0 | 10 | Lake | Browns Bay | 0.62 | 0.73 |
| Campbell River |  |  |  |  |  |  |  |  |  |  |
| Marsh | -125.2570 | 50.0387 | 105 | 105 | 0 | 0 | Estuary | Campbell River | 1.43 | 0.68 |
| Campbell River |  |  |  |  |  |  |  |  |  |  |
| Point | -125.2574 | 50.0476 | 173 | 173 | 0 | 0 | Estuary | Campbell River | 0.45 | 0.61 |
| Cecil Lake | -125.5440 | 50.2361 | 100 | 100 | 0 | 10 | Lake | Amor River | 1.57 | 1.08 |
| Cedar Lake | -125.5664 | 50.2029 | 98 | 30 | 29 | 10 | Lake | Amor River | 3.80 | 1.42 |
| Comida Lake | -125.5283 | 50.1502 | 86 | 86 | 0 | 0 | Lake | Mohun River | 3.70 | 1.64 |
| Comida Stream | -125.5215 | 50.1339 | 192 | 192 | 0 | 0 | Stream | Mohun River | 2.70 | 1.40 |
| Cranberry Lake | -125.4553 | 50.0883 | 66 | 66 | 0 | 10 | Lake | Mohun River | 1.45 | 0.73 |
| Echo Lake | -125.4084 | 49.9848 | 68 | 68 | 0 | 10 | Lake | Quinsam River | 3.28 | 1.34 |
| Farewell Lake | -125.5905 | 50.2013 | 216 | 216 | 0 | 0 | Lake | Amor River | 3.25 | 1.64 |
| Farewell Stream | -125.5992 | 50.2058 | 260 | 260 | 0 | 0 | Stream | Amor River | 1.03 | 1.08 |
| Fry Lake | -125.5720 | 50.0244 | 101 | 101 | 0 | 11 | Lake | Campbell River | 1.67 | 1.20 |
| Gosling Lake | -125.5023 | 50.0483 | 147 | 146 | 0 | 0 | Lake | Campbell River | 3.59 | 1.70 |
| Gray Lake | -125.5953 | 50.0553 | 75 | 30 | 27 | 9 | Lake | Campbell River | 4.72 | 1.35 |
| Higgins Lake | -125.5088 | 50.0680 | 81 | 30 | 29 | 10 | Lake | Campbell River | 2.90 | 1.27 |
| Lawson Lake | -125.5790 | 50.0362 | 82 | 30 | 27 | 10 | Lake | Campbell River | 4.28 | 1.85 |
| Little Goose Lake | -125.4890 | 50.1634 | 76 | 31 | 28 | 10 | Lake | Mohun River | 3.68 | 1.37 |
| Little Mud Lake | -125.5505 | 50.2069 | 60 | 30 | 29 | 10 | Lake | Amor River | 2.90 | 1.17 |
| Lower Campbell |  |  |  |  |  |  |  |  |  |  |
| Lake | -125.4760 | 50.0338 | 30 | 30 | 30 | 10 | Lake | Campbell River | 0.87 | 0.86 |
| Lower Stella Lake | -125.5501 | 50.3107 | 104 | 30 | 30 | 10 | Lake | Pye River | 2.75 | 1.26 |
| McCreight Lake | -125.6390 | 50.3295 | 60 | 30 | 28 | 10 | Lake | Amor River | 3.15 | 1.27 |
| McCreight Stream | -125.6457 | 50.3321 | 60 | 30 | 29 | 10 | Stream | Amor River | 2.05 | 1.57 |
| McNair Lake | -125.5744 | 50.2281 | 60 | 30 | 28 | 10 | Lake | Amor River | 2.00 | 1.01 |
| Merril Lake | -125.5568 | 50.0593 | 38 | 30 | 28 | 10 | Lake | Campbell River | 3.87 | 1.60 |
| Mohun Creek | -125.3767 | 50.1113 | 27 | 27 | 21 | 10 | Stream | Mohun River | 0.96 | 0.71 |
| Mohun Lake | -125.4890 | 50.1640 | 60 | 30 | 28 | 10 | Lake | Mohun River | 2.25 | 1.62 |
| Mud Lake | -125.5541 | 50.1871 | 60 | 30 | 28 | 10 | Lake | Amor River | 2.42 | 1.14 |
| Muskeg Lake | -125.5808 | 50.1989 | 104 | 30 | 28 | 10 | Lake | Amor River | 3.73 | 1.10 |
| Ormud Lake | -125.5267 | 50.1795 | 60 | 30 | 28 | 10 | Lake | Amor River | 2.88 | 1.53 |
| Pye Creek | -125.5772 | 50.3122 | 60 | 29 | 27 | 0 | Stream | Pye River | 1.87 | 1.40 |
| Pye Outlet | -125.5191 | 50.3345 | 73 | 73 | 0 | 10 | Estuary | Pye River | 0.92 | 0.60 |
| Roberts Lake | -125.5428 | 50.2158 | 250 | 250 | 0 | 0 | Lake | Amor River | 2.32 | 1.41 |
| Roberts Stream | -125.5681 | 50.2339 | 354 | 345 | 0 | 0 | Stream | Amor River | 1.24 | 1.16 |
| Sayward Estuary | -125.9464 | 50.3769 | 100 | 100 | 0 | 10 | Estuary | Salmon River | 1.10 | 0.85 |
| Snow Lake | -125.5937 | 50.3127 | 17 | 17 | 3 | 9 | Lake | Pye River | 2.41 | 0.80 |
| Stella Lake | -125.5175 | 50.2992 | 60 | 30 | 30 | 11 | Lake | Pye River | 2.20 | 1.48 |
| Upper Campbell |  |  |  |  |  |  |  |  |  |  |
| Lake | -125.5841 | 49.9953 | 105 | 30 | 29 | 10 | Lake | Campbell River | 1.00 | 0.95 |
| Village Bay Lake | -125.1878 | 50.1708 | 81 | 60 | 59 | 0 | Lake | Village Bay | 3.10 | 1.64 |
| Village Bay Stream | -125.2064 | 50.1675 | 60 | 30 | 29 | 0 | Stream | Village Bay | 1.50 | 1.08 |

**Supplementary Table S2.** NMDS loadings for stickleback diet.

| Prey item | NMDS1 | NMDS2 | NMDS3 | NMDS4 |
| --- | --- | --- | --- | --- |
| Ant | 0.002 | 0.000 | 0.000 | -0.002 |
| Arachnid | 0.003 | 0.003 | -0.004 | -0.003 |
| Bosmina | 0.148 | 0.017 | 0.054 | 0.165 |
| Calanoid | 0.072 | 0.056 | -0.036 | 0.255 |
| Ceratopogonid pupae | 0.140 | 0.129 | -0.142 | 0.851 |
| Ceratopogonid larvae | -0.553 | 0.587 | -0.563 | -0.053 |
| Chironomid pupae | -0.029 | -0.025 | -0.063 | -0.133 |
| Chironomid larvae | -0.637 | -0.714 | -0.118 | 0.194 |
| Chydorus | 0.000 | 0.038 | 0.038 | -0.052 |
| Clams | -0.014 | -0.030 | 0.012 | -0.008 |
| Coleoptera | 0.000 | 0.003 | -0.004 | -0.004 |
| Collembola | 0.005 | 0.006 | 0.006 | -0.001 |
| Cyclopoid | 0.037 | -0.016 | -0.011 | -0.108 |
| Daphnia | -0.041 | -0.026 | 0.046 | 0.027 |
| Diaphnasoma | 0.017 | 0.016 | -0.008 | 0.007 |
| Diptera pupae | 0.028 | 0.009 | 0.000 | -0.050 |
| Diptera adult | 0.005 | 0.023 | -0.012 | -0.128 |
| Diptera larvae |  |  |  |  |
| unidentified | 0.080 | 0.027 | 0.055 | -0.099 |
| Empididae larvae | -0.009 | -0.011 | -0.015 | -0.003 |
| Ephemeroptera | -0.072 | 0.003 | 0.178 | -0.022 |
| Gammarus | -0.464 | 0.342 | 0.772 | 0.153 |
| Harpactacoid | 0.088 | 0.000 | -0.052 | 0.091 |
| Hemiptera | 0.002 | 0.000 | 0.000 | -0.002 |
| Holopedium | 0.000 | 0.001 | -0.001 | 0.000 |
| Hydracarina | 0.000 | -0.014 | -0.011 | -0.002 |
| Leptodoridae | 0.005 | 0.001 | 0.006 | -0.005 |
| Mussel | -0.007 | -0.005 | 0.006 | -0.001 |
| Odonata | 0.002 | 0.000 | 0.000 | -0.002 |
| Ostracod | -0.029 | -0.001 | 0.006 | -0.113 |
| Plecoptera larvae | 0.002 | -0.001 | 0.029 | -0.055 |
| Plecoptera adult | -0.003 | 0.001 | 0.001 | -0.001 |
| Polyphemus | 0.002 | 0.000 | 0.000 | -0.003 |
| Snail | 0.018 | -0.001 | -0.005 | -0.042 |
| Stickle egg | -0.011 | 0.041 | -0.017 | -0.178 |
| Stickle larvae | -0.005 | 0.005 | 0.003 | -0.010 |
| Tabanidae larvae | 0.000 | 0.001 | 0.002 | 0.001 |
| Tanyderidae larvae | -0.001 | 0.006 | -0.007 | -0.008 |
| Tipulid pupae | -0.001 | -0.004 | -0.001 | -0.001 |
| Tipulid larvae | 0.000 | 0.007 | -0.001 | -0.004 |
| Trichoptera pupae | 0.002 | 0.001 | 0.005 | -0.003 |
| Trichoptera larvae | 0.003 | -0.012 | 0.048 | -0.025 |
| Trichoptera adult | 0.002 | 0.006 | -0.008 | 0.002 |
| Percent variance | 7.2 | 7.1 | 6.2 | 5.4 |

**Table S3. Summary of statistical results from the models listed in Table 1 <still in prep>**

**Table S4.** Path analysis results for Figure 3. Statistically significant effects are indicated in bold. Lakes (N = 33) represent the level of replication in this analysis. r^2^ values for variables are 0.473 (parasite richness), 0.686 (benthic diet), 0.306 (standard length), and 0.449 (heterozygosity).

| Variable 1 | Variable 2 | Partial correlation | Std. error | Z | P |
| --- | --- | --- | --- | --- | --- |
| Mean heterozygosity | Mean per-fish parasite richness | **0.360** | **0.174** | **2.07** | **0.038** |
| Benthic diet | Mean per-fish parasite richness | **0.570** | **0.230** | **2.48** | **0.013** |
| Standard length | Mean per-fish parasite richness | 0.075 | 0.159 | 0.47 | 0.635 |
| Elevation | Mean per-fish parasite richness | 0.245 | 0.218 | 1.12 | 0.261 |
| Log lake surface area | Mean per-fish parasite richness | 0.053 | 0.236 | 0.225 | 0.822 |
| Distance from ocean | Mean per-fish parasite richness | 0.244 | 0.219 | 1.12 | 0.264 |
| **Log lake surface area** | **Benthic diet** | **-0.777** | **0.100** | **-7.76** | **<0.001** |
| Standard length | Benthic diet | 0.159 | 0.100 | 1.16 | 0.111 |
| **Log lake surface area** | **Standard length** | **-0.337** | **0.148** | **-2.27** | **0.023** |
| **Distance from ocean** | **Standard length** | **0.516** | **0.148** | **3.48** | **0.001** |
| **Log lake surface area** | **Mean heterozygosity** | **0.366** | **0.133** | **2.54** | **0.011** |
| **Distance from ocean** | **Mean heterozygosity** | **0.553** | **0.180** | **3.07** | **0.002** |
| **Elevation** | **Mean heterozygosity** | **-0.741** | **0.177** | **-4.20** | **<0.001** |

